# DEBKS: A Tool to Detect Differentially Expressed Circular RNA

**DOI:** 10.1101/2020.10.14.336982

**Authors:** Zelin Liu, Huiru Ding, Jianqi She, Chunhua Chen, Weiguang Zhang, Ence Yang

## Abstract

Circular RNAs (circRNAs) are involved in various biological processes and in disease pathogenesis. However, only a small number of functional circRNAs have been identified among hundreds of thousands of circRNA species, partly because most current methods are based on circular junction counts and overlook the fact that circRNA is formed from the host gene by back-splicing (BS). To distinguish between expression originating from BS and that from the host gene, we present DEBKS, a software program to streamline the discovery of differential BS between two rRNA-depleted RNA sequencing (RNA-seq) sample groups. By applying real and simulated data and employing RT-qPCR for validation, we demonstrate that DEBKS is efficient and accurate in detecting circRNAs with differential BS events between paired and unpaired sample groups. DEBKS is available at https://github.com/yangence/DEBKS as open-source software.

## Introduction

Circular RNAs (circRNAs) are a class of long noncoding RNAs with a covalently closed continuous loop structure that is formed via back-splicing (BS). Although circRNAs are believed to play important regulatory roles in a number of biological processes [1–4], only a small number of functional circRNAs have been recognized among hundreds of thousands of circRNA species [5–8]. A straightforward strategy to identify functional circRNAs is to detect differentially expressed circRNAs (DECs) that may contribute to certain traits or diseases [9–11]. However, the identification of DECs is still restricted by software limitations.

Previously developed software, such as find_circ [12], CIRI2 [13], and CIRCexplorer2 [14], focused on the identification of circRNAs, as well as quantification by circular junction counts (CJCs), while recent tools, including CIRI-full [15], CircSplice [16], and CircAST [17], could detect circRNA alternative splicing and quantify CJCs at the isoform level. As these programs are not able to detect DECs, differentially expressed gene (DEG) detection software (mostly DESeq2 [18] and edgeR [19]) is employed to analyze the data obtained with these CJC-based quantification tools. However, the circRNA expression level is usually associated with the host gene [20], and it is therefore difficult to differentiate whether the difference is due to BS or is only a byproduct of the host gene.

Some programs, such as DCC [21], CIRI2, CIRIquant [22], and CLEAR [23], could measure circRNA expression level by the ratio of BS, instead of CJCs, to normalize the noise of host gene expression. Given that the BS ratio is not applicable for DESeq2 and edgeR, here we report **DE b**ac**k**-**s**plicing (DEBKS), a tool that detects differential BS between sample groups with paired or unpaired rRNA-depleted RNA-seq data. DEBKS includes four functional modules: *merge* for reformatting junction counts from circRNA detection software, *count* for quantifying expression level of circRNA linear counterpart, *anno* mainly for estimating circRNA length, and *dec* for detecting DECs. Using public, in-house, and simulated RNA-seq datasets, we demonstrate that DEBKS effectively improves the ratio of BS calculation and has a better performance in DEC detection than do current methods.

## Method

### Workflow of DEBKS for identifying differentially expressed circRNAs

DEBKS is designed to analyze DECs from rRNA-depleted RNA-seq without erasing linear RNA. DEBKS includes four modules: *merge, count, anno*, and *dec* (**Figure 1A**). After detecting circRNAs by users, *merge* can collect and reformat CJCs from files generated using detection software. This module can also simultaneously collect linear junction counts (LJCs), if available, which is defined as junction counts across flanking introns of candidate back-spliced sites. Alternatively, *count* is employed for calculating LJCs from BAM files produced by RNA-seq aligners, such as HISAT2 [24] and STAR [25]. Similar to the percentage spliced in [26], the ratio of BS events is measured by percentage back-spliced in (PBSI, *ψ*), which is estimated by

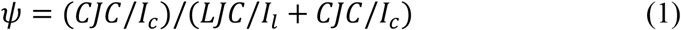

where *I_l_* represents the effective linear length and *I_c_* represents the effective back-spliced length (**Figure 1B**). The values of *I_l_* and *I_c_* are estimated by

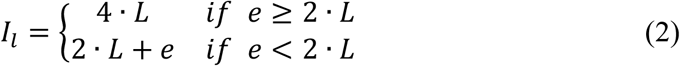

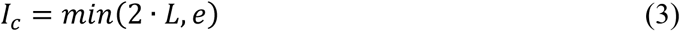

where *L* represents the RNA-seq read length minus the minimum overhang length, and *e* represents the circRNA length, which can be optionally predicated with gene annotation by *anno* or full-length analysis software, such as CIRI-full and CircAST. For smaller circRNAs (*e* <2 · *L*), *ψ* is adjusted with smaller effective lengths. For larger circRNAs (*e* ≥ 2 · *L*), *ψ* is equivalent to the calculation of the ratio in other programs, such as CIRI2 and CIRIquant. Finally, the statistical framework of rMATS [26], which was developed for detecting differential alternative splicing (AS), is employed to compare the difference of *ψ* (|Δ*ψ*|) with a user-defined threshold between the paired or unpaired sample groups.

**Figure 1.**
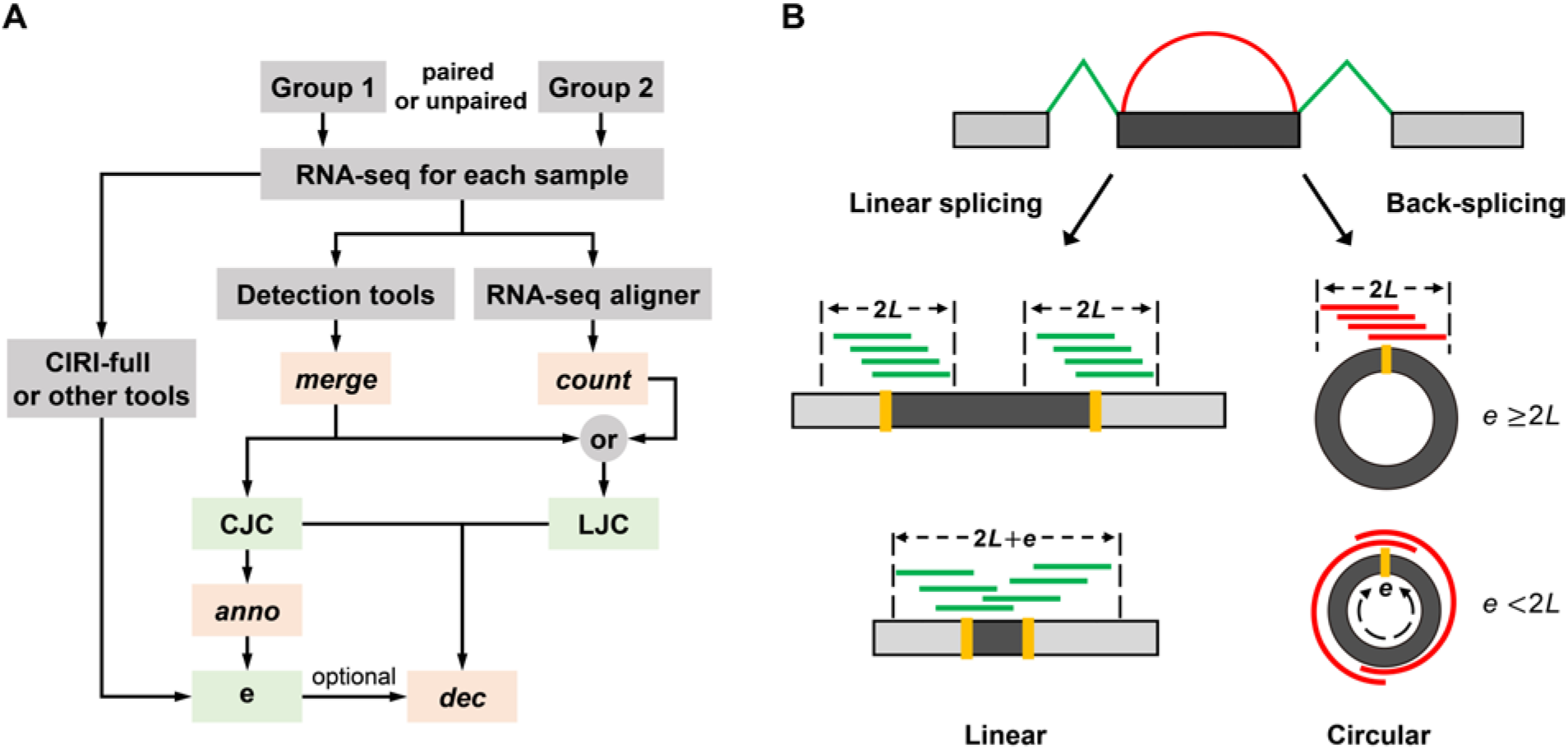
DEBKS workflow. (A) The workflow of DEBKS consists of four modules: *merge, count, anno*, and *dec*. (B) The schematic diagram of back-splicing. Red reads represent CJCs. Green reads represent LJCs. CJCs, circular junction counts; LJCs, linear junction counts; L, RNA-seq read length minus the minimum overhang length; e, circRNA length.

### Public RNA-seq datasets

Two public rRNA-depleted RNA-seq datasets (SRP050149 and SRP156355) from the NCBI sequence read archive (SRA) were employed to verify the performance of DEBKS. SRP050149 includes four samples of HEK293 cells with the codepletion of ADAR1 and ADAR2 by RNAi (ADAR1,2 knockdown) and four untreated samples. SRP156355 includes five invasive ductal carcinoma (IDC) samples and five paired normal samples.

### Mouse model of transient middle cerebral artery occlusion

Transient focal ischemia of three 6- to 7-week-old male mice (C57BL/6) weighing 21 ± 1 g was induced by middle cerebral artery occlusion (MCAO) using the endovascular suture method [27]. The operation was performed under isoflurane anesthesia. The right external carotid artery (ECA), internal carotid artery (ICA) and common carotid artery (CCA) were exposed under a stereoscopic anatomical microscope. The suture was gently pushed from the ECA to the ICA to block the origin of the MCA (9-10 mm). After 1 hour, the suture was withdrawn to allow MCA reperfusion. After 6 hours, mice were sacrificed, and brain tissue from the ischemic core area on the right side of the brain was removed, while normal brain tissue from the corresponding area on the left side was removed and utilized as a control. Next, RNA extraction was performed immediately.

### RNA extraction

Total RNA of the ischemic core and normal areas were extracted by the FastPure Cell/Tissue Total RNA Isolation Kit (RC101, Vazyme Biotech Co., Ltd., Nanjing, Jiangsu, China) with 10-20 mg of brain tissue according to the manufacturer’s instructions.

### Library preparation for RNA-seq

The library construction and sequencing of rRNA-depleted RNA was performed by Annoroad Gene Technology (Beijing, China). A total of 3 μg RNA per sample was used as the input material for rRNA removal with the Ribo-Zero rRNA Removal Kit (MRZG12324, Illumina, San Diego, CA, USA). Libraries were generated using the NEBNext^®^ Ultra™ Directional RNA Library Prep Kit for Illumina^®^ (E7420, NEB, Ipswich, MA, USA) after sequencing on an Illumina HiSeq X platform with 150-bp paired-end reads.

### RT-qPCR analysis

Total RNA (500 ng) was reverse-transcribed into complementary DNA (cDNA) by using HiScript^®^III RT Strand SuperMix for qPCR (+gDNA wiper) (R323, Vazyme Biotech Co., Ltd., Nanjing, Jiangsu, China). For each circRNA, two primers inside BS and two primers outside BS were designed to detect circular junctions and upstream and downstream linear junctions (Figure S1, Table S1). RT-qPCR was performed by using ChamQ™ Universal SYBR^®^ qPCR Master Mix (Q711, Vazyme Biotech Co., Ltd, Nanjing, Jiangsu, China) on an ABI7500 system (Applied Biosystems, Foster, CA, USA) according to the manufacturer’s procedures and with β-actin as an internal reference gene. Then, *ψ* is calculated by determining the ratio of the relative expression level of the circular junction to the sum of the circular junctions and the average of the upstream and downstream linear junctions.

### Identification of DECs and DEGs in real RNA-seq

For each RNA-seq, raw reads were mapped to the reference genome (hg19 for SRP050149 and SRP156355 and mm10 for mouse RNA-seq) by STAR (v2.5.3a). Next, the LJCs of circRNAs were calculated by DEBKS, while CJCs of circRNAs and counts of host genes were calculated by CIRCexplorer2 [14] (v2.3.8) and featureCounts (v1.6.0) [28], respectively. Only circRNAs with average CJCs ≥ 2 were kept in the following DE analysis. DECs and DEGs between the two groups were analyzed using DEBKS and DESeq2, respectively.

### Simulation of rRNA-depleted RNA-seq

For the simulated RNA-seq dataset1, we randomly selected 10,000 circRNA host genes based on human gene annotation (GENCODE v19) for 3 vs. 3 sample groups. We assumed that each host gene contained only one linear product but that half genes contained one circular product, while the other half contained two circular isoforms (same back-spliced site with exon skipping in one isoform). For each gene, the total number of linear and circular RNA molecules was sampled from a normal distribution, with the mean and the standard deviation (SD) being derived from the sum of CJCs and LJCs of the in-house MCAO mouse RNA-seq. For the two sample groups, ratios of BS were simulated with 9500 pairs from a beta distribution (p = 1, q = 12) and 500 pairs from a beta distribution (p = 7, q = 10). Half of the ratio pairs had differences greater than 5% as DEC, and the ratio of BS for each sample was simulated with the mean equal to the group ratio and the SD at five different levels, that is, 0.01, 0.02, 0.05, 0.10, and 0.20. Next, the number of circular and linear molecules was calculated with the total number of molecules and the ratio of BS. For circRNAs with AS, the percentages of one circRNA isoform against total circular molecules were simulated with uniform distribution. Next, the number of each circular isoform molecule was calculated with the percentage and the total number of circular molecules. Finally, simulated rRNA-depleted RNA-seq was generated with simulated numbers of circular and linear molecules.

For the simulated RNA-seq dataset2, again, 10,000 circRNA host genes based on human gene annotation (GENCODE v19) were randomly selected for 3 vs. 3 sample groups. Each gene contained one linear product and two circular isoforms from the same back-spliced site. The total number of linear and circular RNA molecules was sampled as described in dataset1. The ratio pairs between groups of each circular isoform against the circular and the linear isoforms were simulated with beta distribution (p = 1, q = 12 for 9500 ratio pairs; p = 7, q = 10 for 500 ratio pairs). For the two circular isoforms from the same gene, there was one and only one isoform that had a difference of more than 5% from DEC at the isoform level. Next, the number of circular and linear molecules was calculated with the total number of molecules and the ratios of BS. Finally, simulated rRNA-depleted RNA-seq was generated with a number of circular and linear molecules.

### Identification of DECs in simulated RNA-seq

For each RNA-seq, raw reads were mapped to the reference genome (hg19) by BWA mem [29] (v0.7.17-r1188). circRNAs were detected by CIRI2 (v2.0.6) with gene annotation (GENCODE v19). In the simulated RNA-seq dataset1, DEBKS, CircTest [21], and Fisher’s exact test were employed to identify DECs based on the CJCs and LJCs from the output file of CIRI2. In the simulated RNA-seq dataset2, the CJCs of each isoform of circRNAs were detected by CIRI-full, and the corresponding LJCs were obtained from CIRI2.

### Ethics statement

All experimental animals were purchased from the Experimental Animal Center of Peking University. All protocols involving animals, behavioral testing, surgery and animal care were approved by the laboratory animal welfare ethics committee.

### Data availability

RNA-seq data of the mouse MCAO model have been deposited in the GSA:CRA002696. The source code of DEBKS and the script to simulate the RNA-seq dataset are available at https://github.com/yangence/DEBKS.

## Results

### Identification of DECs with DEBKS in two public RNA-seq datasets

We first evaluated the performance of DEBKS using two public rRNA-depleted RNA-seq datasets. The first dataset includes samples of HEK293 cells with or without the knockdown of ADAR1,2. Given that RNA editing is exclusive to circRNA formation [9], we expected to observe more upregulated circRNAs in the ADAR1,2 knockdown treatment. With |Δ*ψ*| > 0.01 at FDR < 0.05, DEBKS detected 14 DECs, all of which were upregulated in ADAR1,2 knockdown cells as expected (**Figure 2A**).

**Figure 2.**
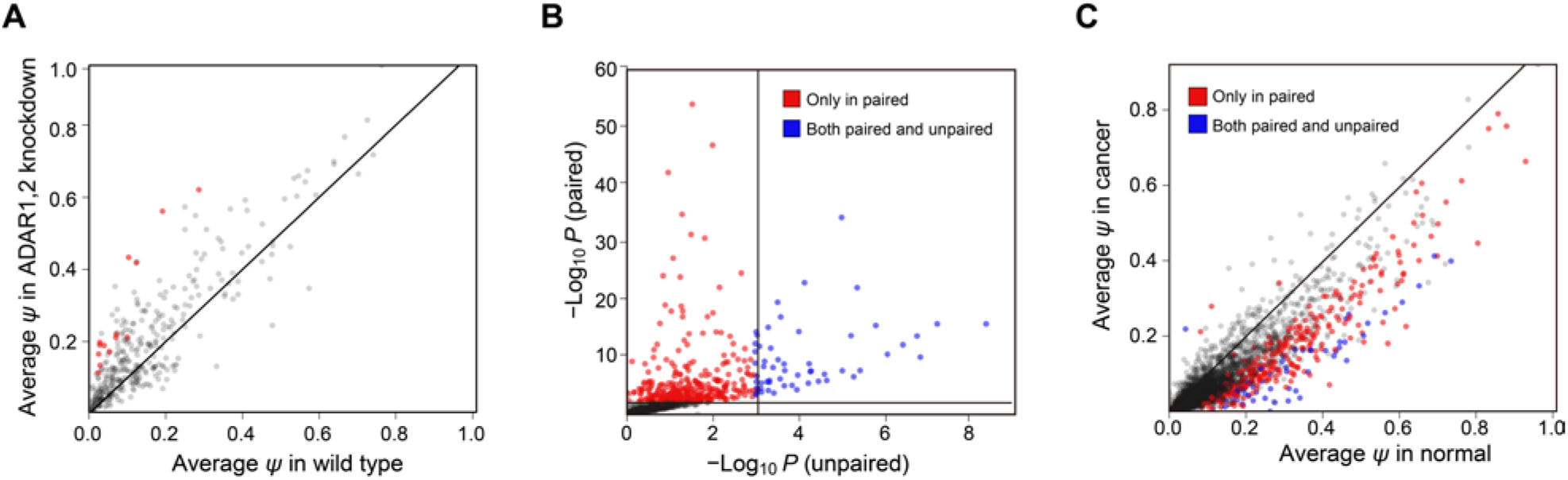
Distribution of DECs in public RNA-seq data. (A) Average *ψ* in ADAR1,2 knockdown and normal HEK293 groups. Red points represent DECs detected by DEBKS. (B, C) *P*-values (B) and average *ψ* (C) of circRNAs detected by DEBKS with paired or unpaired model between IDC and paired normal. Red points represent DECs only detected in paired model, and blue points represent DECs detected in both paired and unpaired models. DECs, differentially expressed circRNAs.

The second dataset includes 10 RNA-seq of IDC and paired normal samples. With or without the parameter ‘-p’, DEBKS detected 377 DECs with the paired model and detected 62 DECs with the unpaired model (**Figure 2B**). Most (372 of 377) DECs detected by the paired model were downregulated in cancer samples, which is in keeping with the results of a previous study [30]. The |Δ*ψ*| of 315 DECs only detected in the paired model was significantly (*P* = 2.6×10^−5^, as determined by a two-tailed Student’s t-test, **Figure 2C**) lower than the 62 DECs detected by both models, indicating that the statistical framework with paired information can result in considerably improved sensitivity in paired samples.

### DEBKS was consistent with RT-qPCR validation in mouse brain

Next, we detected DECs between the ischemic core and control area of the mouse brain with MCAO, which can induce differential expression of circRNAs in the ischemic core [31]. At the threshold of |Δ*ψ*| > 5% and FDR < 0.05, a total of 74 DECs (10 upregulated and 64 downregulated in ischemic area) were identified with rRNA-depleted RNA-seq by DEBKS with a paired statistical model. Among highly expressed (average CJC ≥ 10) DECs, five were randomly selected for validation by RT-qPCR, and they were all consistent with RNA-seq (**Figure 3A, B**). Notably, we did not detect any DECs with either DEBKS in the unpaired model or CircTest of DCC, suggesting that paired samples are more powerful for detecting DECs, especially with DEBKS, the only software demonstrated to detect differential BS for paired groups with replicates to date.

**Figure 3.**
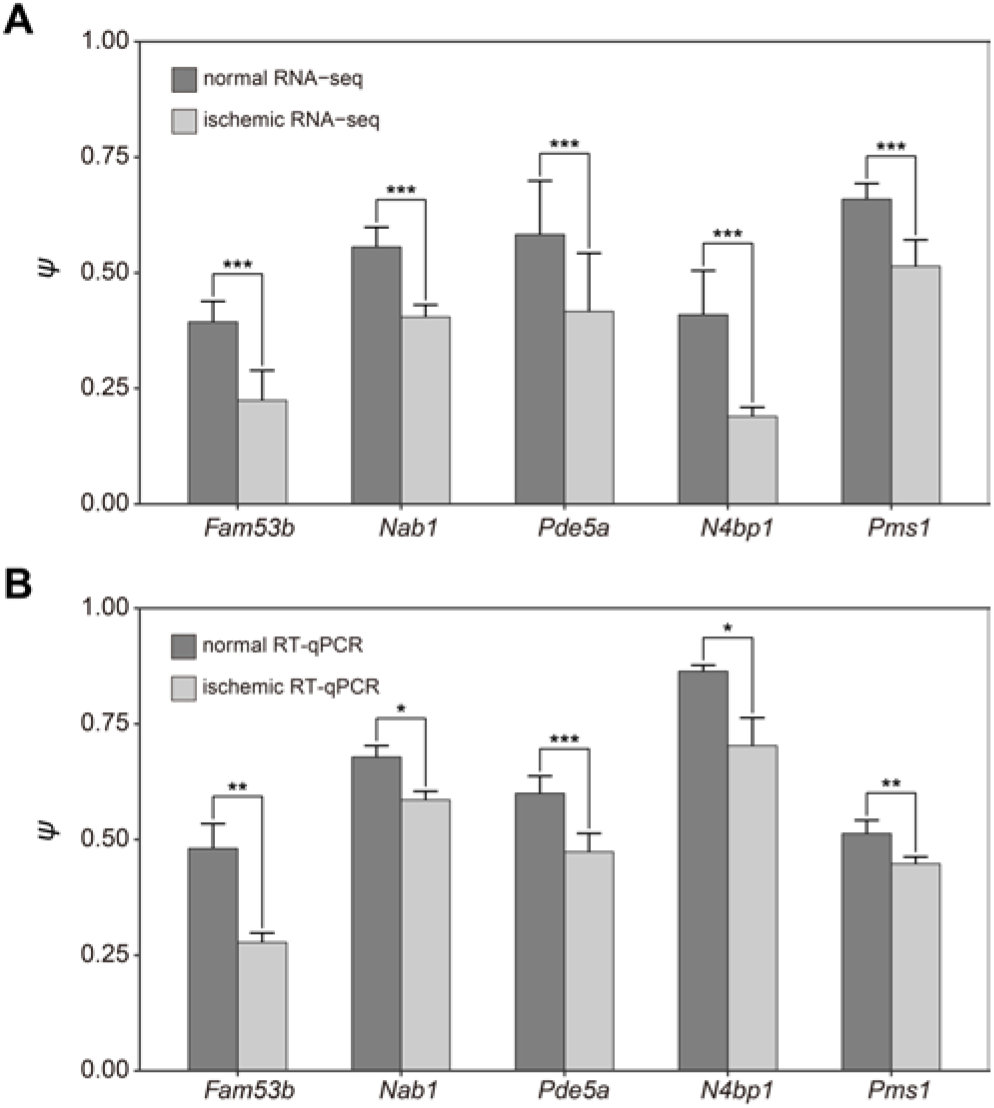
Five circRNAs validated by RT-qPCR in the mouse brain. *ψ* of five circRNAs in ischemic core and control areas of the mouse brain with RNA-seq (A) and RT-qPCR (B). *P*-value is evaluated by DEBKS for RNA-seq, and single tailed paired Student’s t-test for RT-qPCR. *, **, *** represent *P* < 0.1, *P* < 0.05, and *P* < 0.01.

### DEBKS performed better than current methods in simulation data

To further evaluate the performance of DEBKS, we simulated rRNA-depleted RNA-seq based on the real junction count distribution with five intragroup SDs, that is, 0.01, 0.02, 0.05, 0.10, and 0.20, which represented the variability of the ratio of BS among replicates. In the simulated data, the ratio of BS followed the beta distribution, which could better reflect the ratio distribution in real RNA-seq (Figure S2A, B). We first compared the performance of BS ratio quantification with other programs, including CIRI2, CIRIquant, DCC, CIRCexplorer2, CLEAR, and sailfish-cir [32] (**Figure 4**). We found that *ψ* calculated by CJCs from CIRI2 and LJCs from DEBKS was the most consistent with the simulated ratio. In addition, when combined with LJCs from DEBKS, *ψ* was more consistent with simulated data than the ratio derived from CIRI2, CIRIquant, or DCC alone, supporting that DEBKS possessed an advantage for ratio calculation.

**Figure 4.**
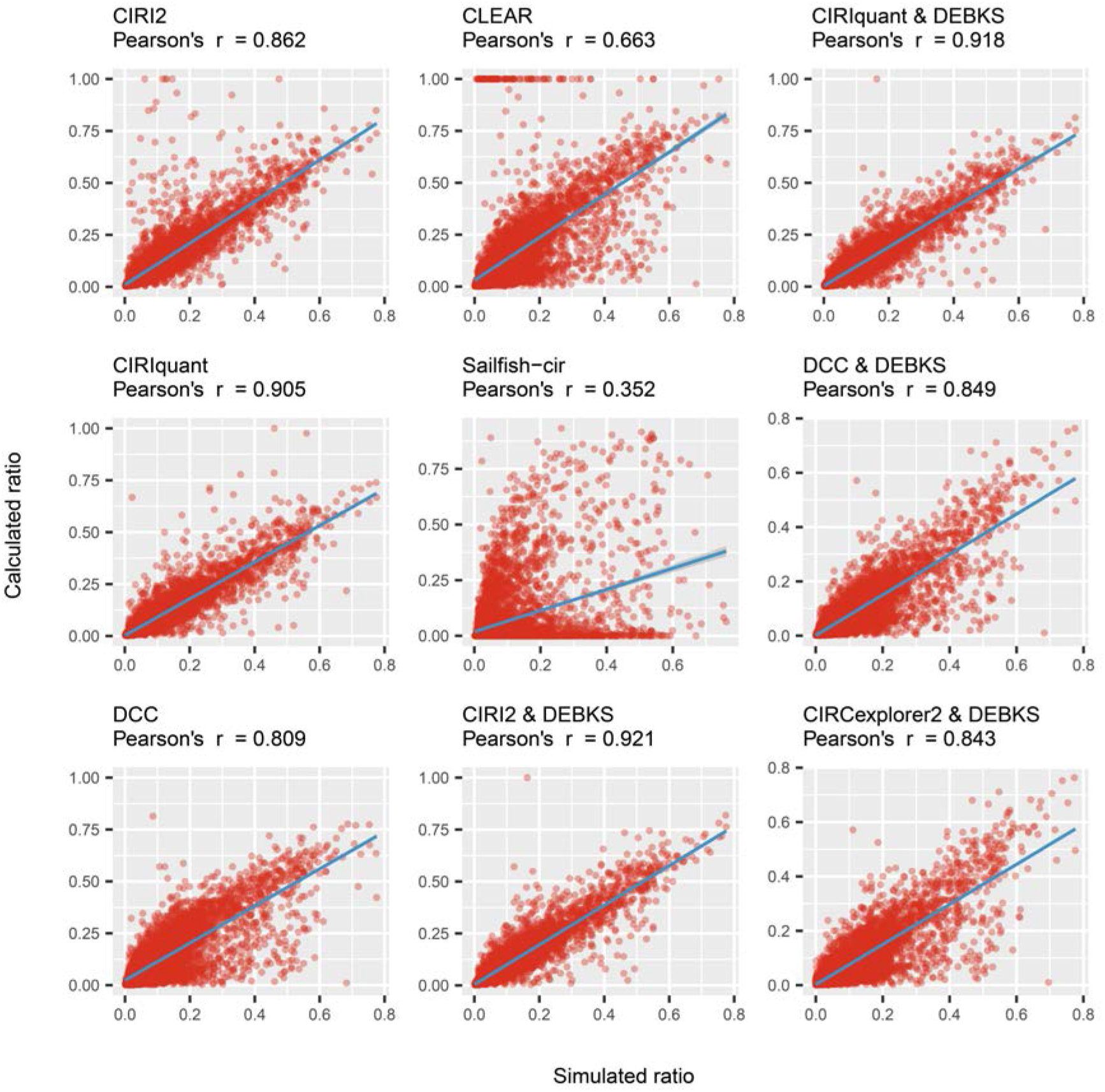
BS ratio calculation with simulated dataset1 is improved in current tools when combining with DEBKS. For CIRI2, CIRIquant, and DCC, BS ratios were calculated by *CJC*/(*LJC*/2 + *CJC*). Because CLEAR and Sailfish-cir used normalized circular (C) and linear (L) expression, we first calculated the ratio by C/(L/2 + C) and *C*/(*L* + *C*) separately and selected *C*/(*L*/2 + *C*) for CLEAR and *C*/(*L* + *C*) for Sailfish-cir based on better Pearson’s *r*. For the *ψ* calculated by DEBKS, CJCs were from CIRI2, CIRIquant, DCC, and CIRCexplorer2, while LJCs were calculated by DEBKS *count* module with STAR alignment. Blue lines indicate the fitting line. BS, back-splicing; CJCs, circular junction counts; LJCs, linear junction counts.

With minimal total CJCs of all samples ranging from 5 to 40 counts and SD from 0.01 to 0.20 in the simulated dataset1, we evaluated the performance of *dec* of DEBKS, as well as CircTest and Fisher’s exact test (FET) for comparison (**Figure 5**). In terms of area under receiver operating characteristic (AUC) values and true positive rate (TPR) at a 5% false positive rate (FPR), DEBKS was determined to perform better than any combination of minimal total CJCs and SD, which indicated that DEBKS was powerful in DEC detection for both weakly and strongly expressed circRNAs. However, by applying CIRI-full to calculate CJCs of each circular isoform in the simulated dataset2, we observed that DEBKS possessed low accuracy for detecting DECs at the isoform level (Figure S3), which was primarily observed because of the inaccurate quantification of CJCs (Figure S4).

**Figure 5.**
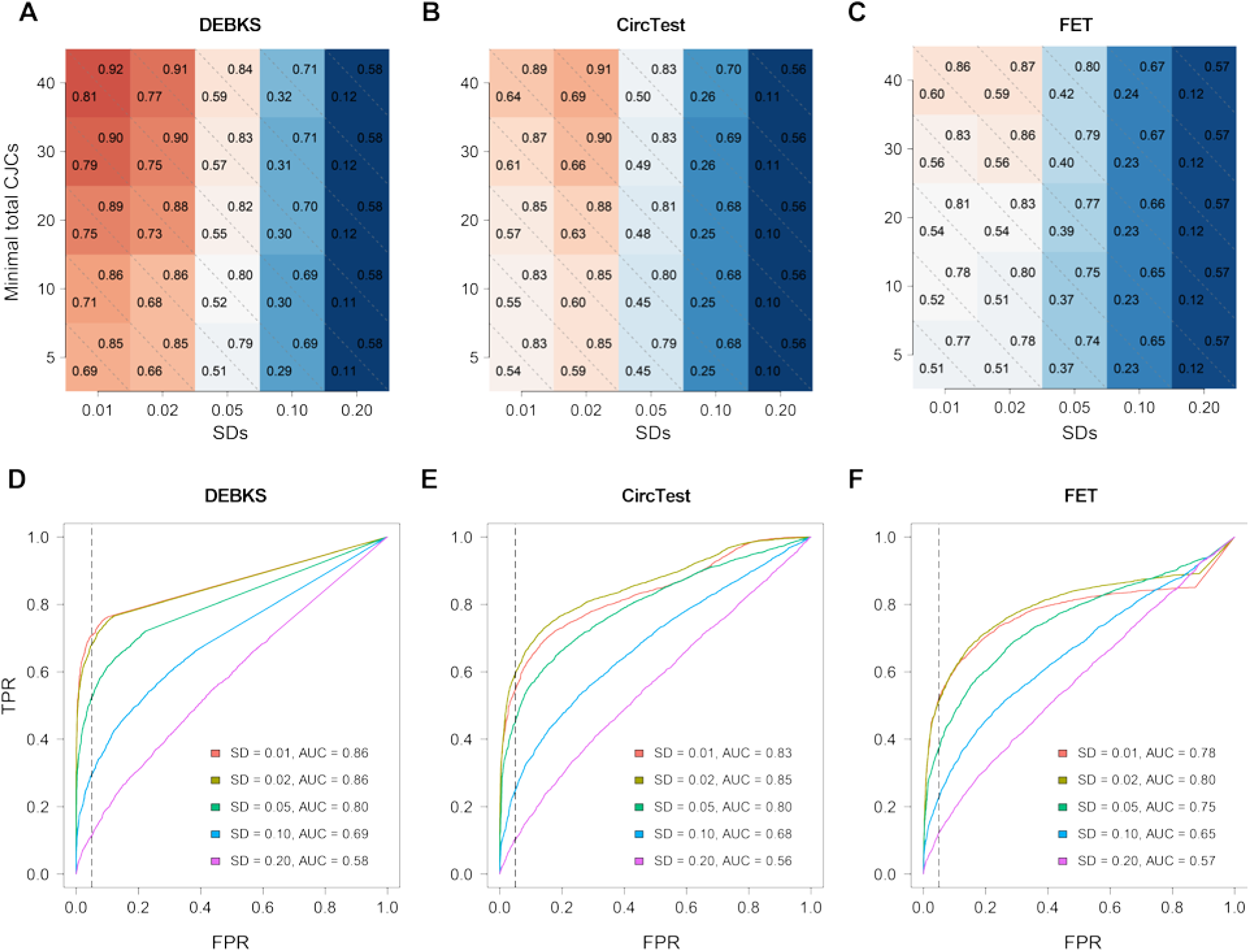
DEBKS shows higher TPR and AUC than CircTest and FET under various group variance and junction counts with the simulation dataset1. (A-C) The values of TPR at 5% FPR (lower triangle) and AUC (upper triangle) of DECs identified by DEBKS (A), CircTest (B), and FET (C) for circRNAs with minimal total CJCs of all samples ranging from 5 to 40 counts and SDs from 0.01 to 0.20. (D-F) ROC curves of five simulation results at SDs from 0.01 to 0.20 for circRNAs with minimal total CJCs of 10 counts for DEBKS (D), CircTest (E), and FET (F). The vertical dashed line indicates an FPR = 0.05. TPR, true positive rate; FPR, false positive rate; AUC, area under receiver operating characteristic; SDs, standard deviations. ROC: receiver operating characteristic; CJCs, circular junction counts.

### DEBKS was less heavily influenced by host gene expression than was DESeq2

Given the low abundance of circRNA in rRNA-depleted RNA-seq, ratio-based methods were considered to be more likely to detect DECs than CJC-based methods, such as DESeq2. By using DESeq2 at FDR < 0.05, we failed to detect any DECs in the knockdown of ADAR1,2 and MCAO RNA-seq and detected 34 DECs in IDC RNA-seq, among which 47% (16/34) were also detected by DEBKS. In addition, the host genes of DECs detected by DESeq2 were considerably more likely to be DEGs (odds ratio = 7.33, *P* = 4.02 × 10^−8^, two-tailed FET) compared to DEBKS with the paired model (odds ratio = 1.08, *P* = 0.67, two-tailed FET) and the unpaired model (odds ratio = 1.63, *P* = 0.15, two-tailed FET), suggesting that DEBKS identified DECs with less influence from host gene expression.

## Discussion

Given that circRNAs are BS isoforms of host genes, we developed DEBKS to identify DECs at the splicing level by BS ratio, which normalizes CJCs with local LJCs. In addition to CJC-based methods, ratio-based methods may help to identify potential functional circRNAs based on the biogenesis mechanism of circRNA [4]. In our evaluation, DEBKS efficiently identified DECs with less influence from host gene expression. For example, among the 3080 circRNAs detected in IDC RNA-seq, a strong correlation of log_2_ (fold change of expression level) (log_2_FC) was observed between circRNA and the host gene (Pearson’s *r* = 0.53, *P* < 2.2 × 10^−16^) (Figure S5), indicating that circRNA expression level was significantly affected by host gene expression. In contrast, only a weak correlation was observed between circRNA Δ*ψ* and log_2_FC of the host gene (Pearson’s *r* = 0.06, *P* = 4.7 × 10^−4^), suggesting that BS events measured by |Δ*ψ*| were less strongly influenced by the expression level of the host gene.

At present, several software, including CircTest and CIRIquant, can detect DECs with ratio-based methods. Compared with simulated data, DEBKS shows better performance for length adjustment of smaller circRNA and employment of the statistical framework of rMATs. In addition, DEBKS could be applied in paired or unpaired groups with replicates, which is distinct from CIRIquant, which utilizes a ratio-based test to detect DECs between groups without replicates.

Because DEBKS is based on the ratio of CJCs and LJCs, it is only applicable in RNA-seq analyses of both circRNA and linear RNA, such as rRNA-depleted or exome capture RNA-seq [10]. In addition, intergenic circRNA without a linear counterpart, which is estimated to account for less than 10% of total circRNA [33], is not suitable for DEBKS.

## CRediT statement

**Liu Zelin:** Conceptualization, Software, Formal Analysis, Writing - Original Draft, Writing - Review & Editing, Visualization. **Ding Huiru:** Methodology, Validation, Writing - Review & Editing. **She Jianqi:** Investigation, Data curation, Writing - Original Draft. **Chen Chunhua:** Resources, Project administration, Funding acquisition. **Zhang Weiguang:** Supervision, Resources, Writing - Review & Editing. **Yang Ence:** Conceptualization, Supervision, Resources, Funding acquisition, Writing - Original Draft, Writing - Review & Editing.

## Acknowledgements

This study was supported by Beijing Municipal Science and Technology Commission, China (Z181100001518005) and Chinese Institute for Brain Research, Beijing, China to EY and National Natural Science Foundation of China (81873769) to CC.

## Supplementary material

**Figure S1.**
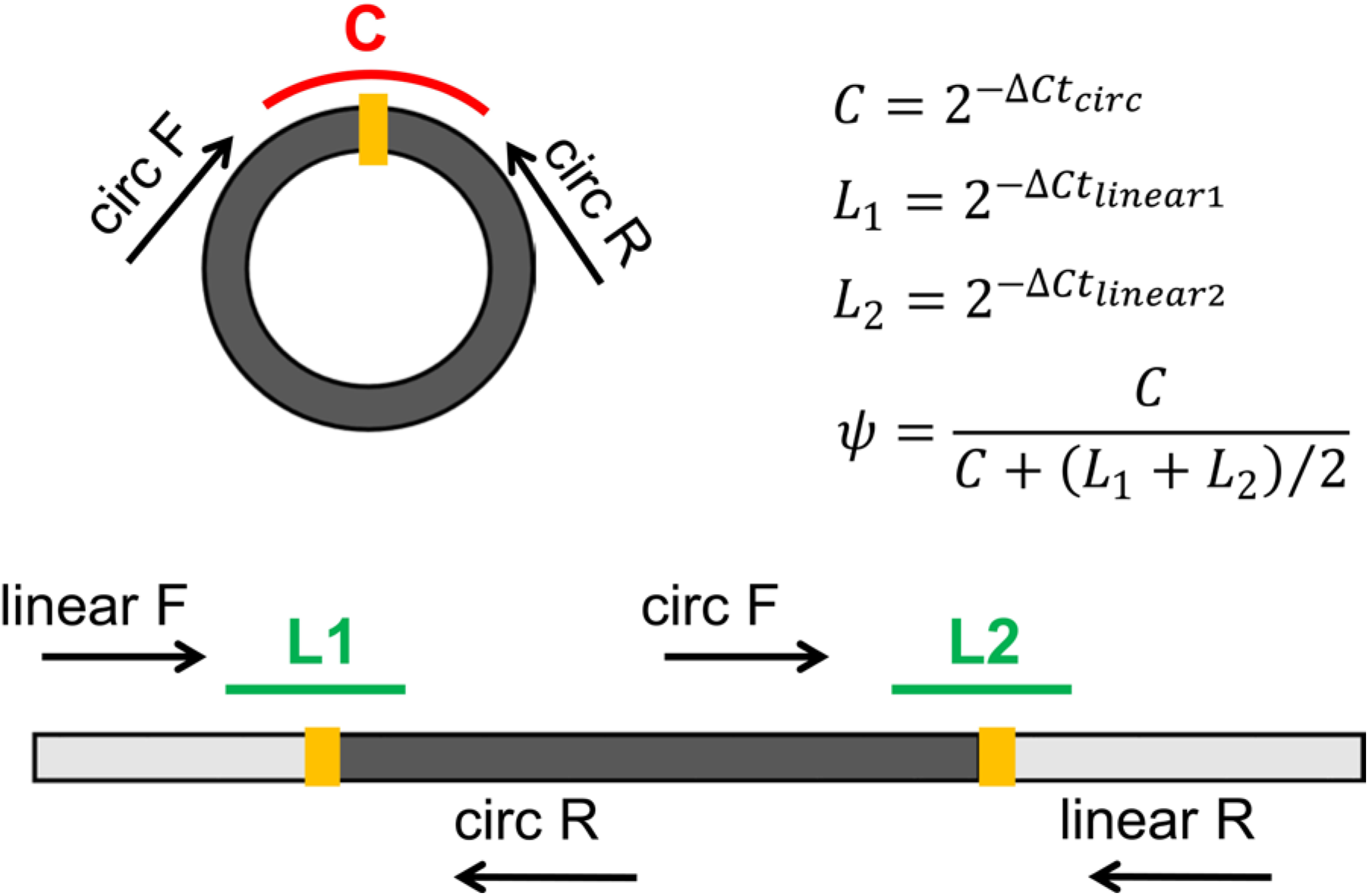
Schematic diagram of primers for RT-qPCR. circF and circR are designed for C; linearF and circR are designed for L1; linearR and circF are designed for L2.

**Figure S2.**
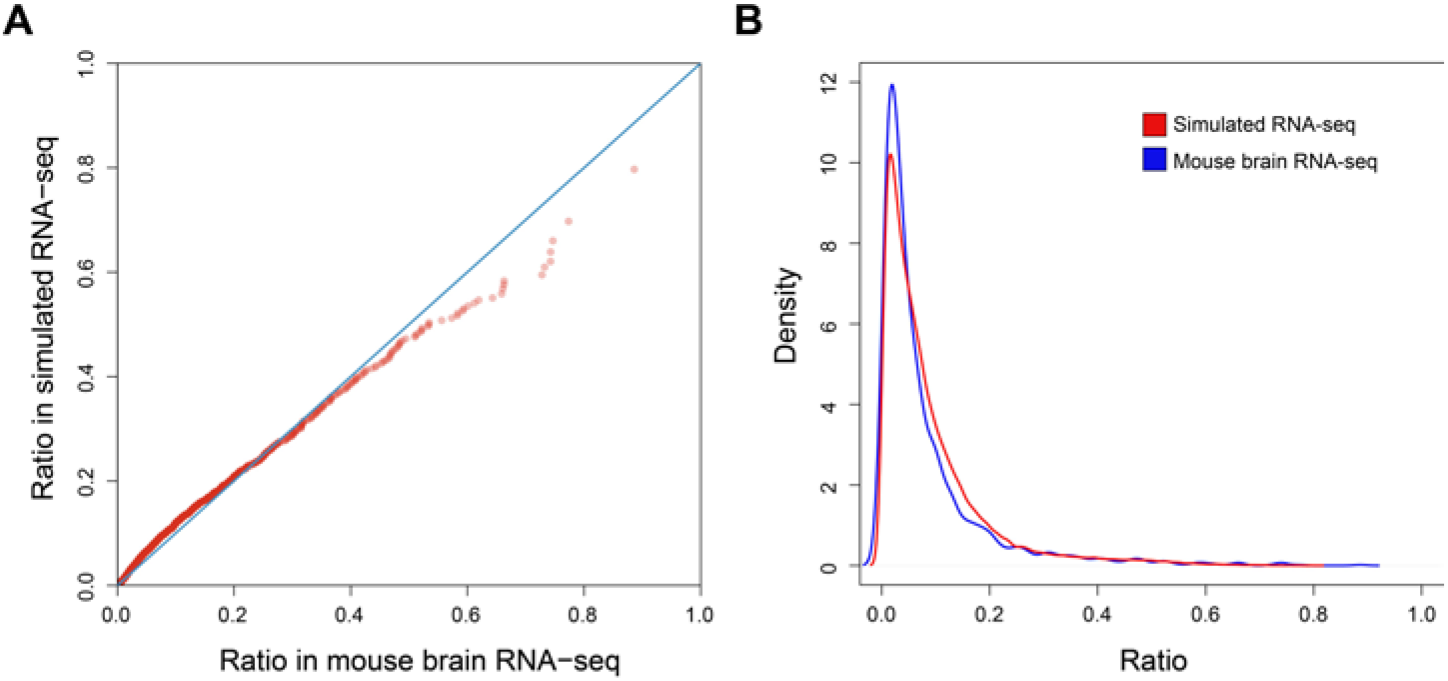
Simulated RNA-seq shows similar distribution with real data. QQplot (A) and density distribution (B) of ratio of BS in simulated RNA-seq and mouse RNA-seq. BS, back-splicing.

**Figure S3.**
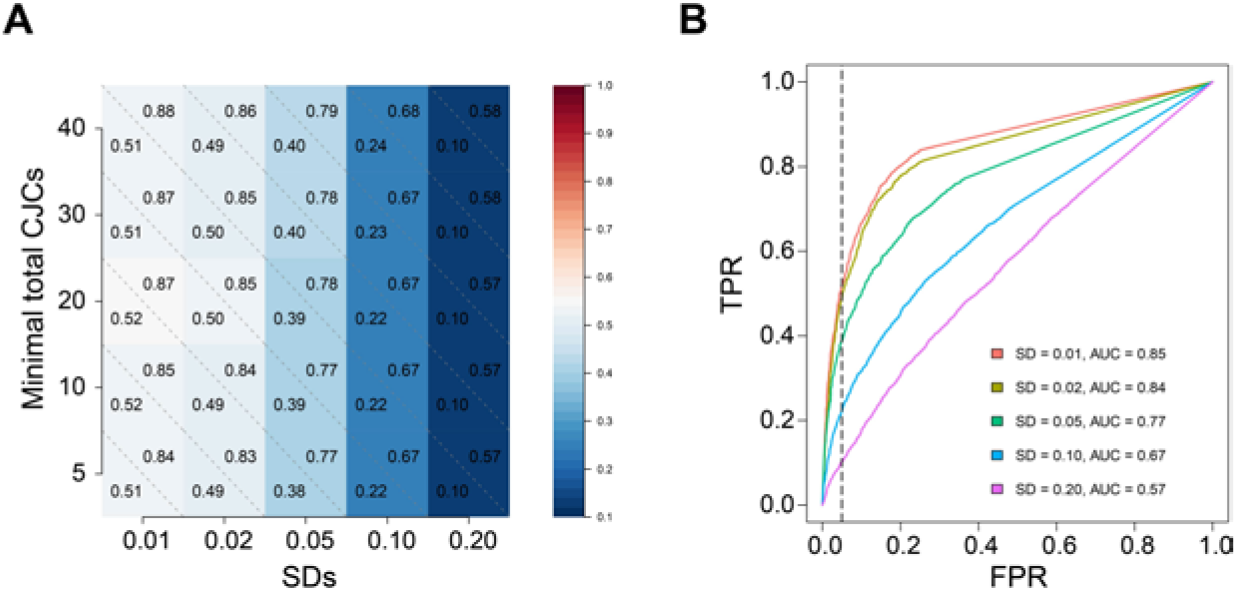
Evaluation of the influence of group variance and junction counts on detection accuracy for differential alternative splicing in circRNA with the simulation dataset2. (A) The values of TPR at 5% FPR (lower triangle) and AUC (upper triangle) of DECs identified by DEBKS for minimal total CJCs of all samples ranging from 5 to 40 counts and SDs from 0.01 to 0.20. (B) ROC curve of five simulation results at SDs from 0.01 to 0.20 for circRNAs with minimal total CJCs of 10 counts. The vertical dashed line indicates an FPR = 0.05. TPR, true positive rate; FPR, false positive rate; AUC, area under receiver operating characteristic; SDs, standard deviations. ROC: receiver operating characteristic; CJCs, circular junction counts.

**Figure S4.**
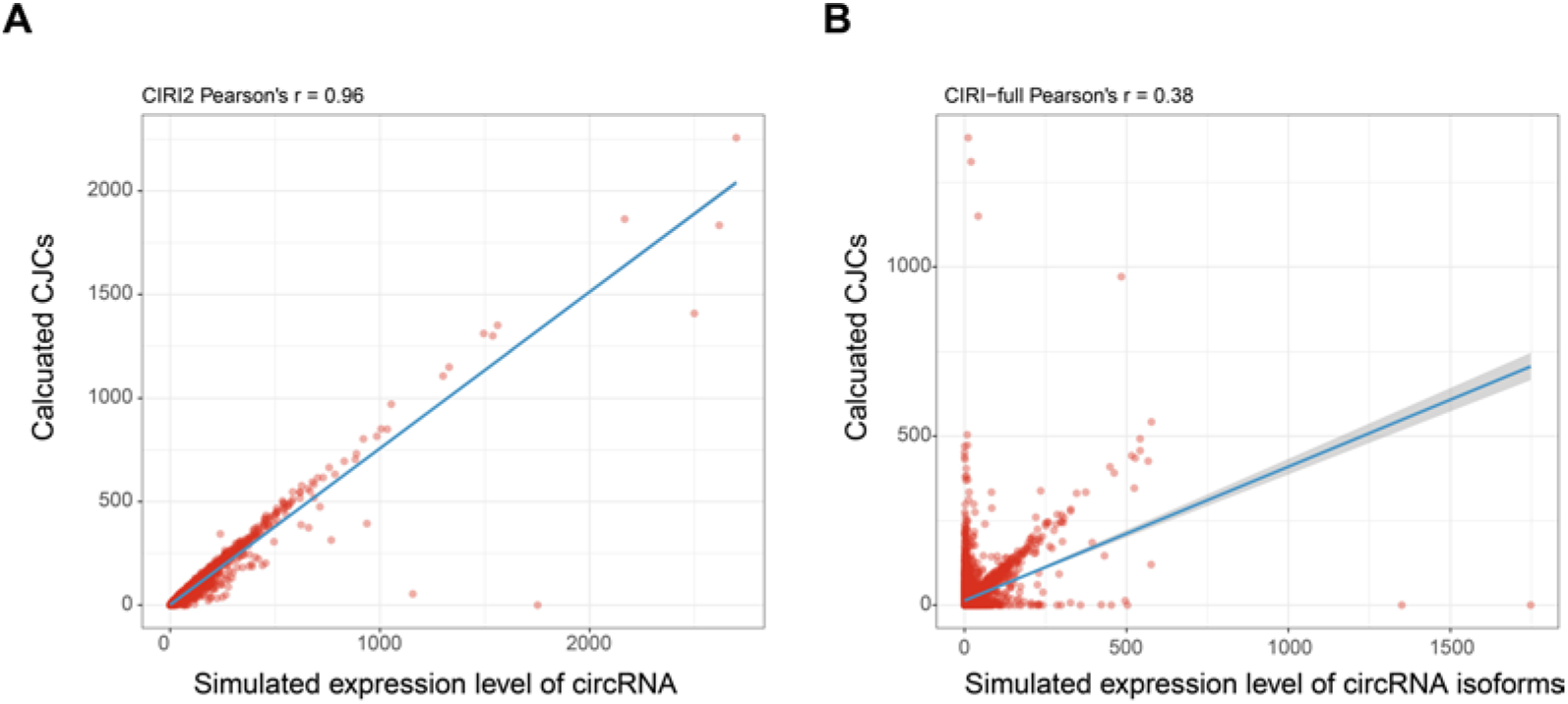
CJCs calculated at isoform level is less accurate than total CJCs. Plot of calculated CJCs from CIRI2 (A) and CIRI-full (B) based on simulated RNA-seq dataset2 with simulated expression level of circRNA and circRNA isoforms. CJCs, circular junction counts.

**Figure S5.**
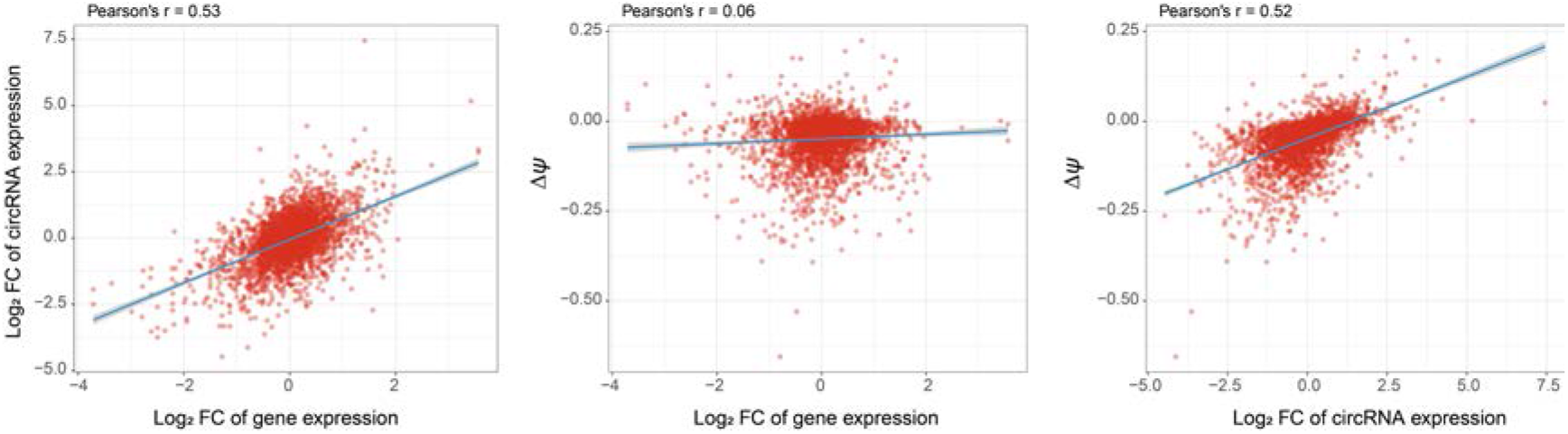
Δ*ψ* is less influenced by fold change of gene expression. Correlations among fold change of gene expression, fold change of circRNA expression, and Δ*ψ* in IDC RNA-seq.

**Table S1.**
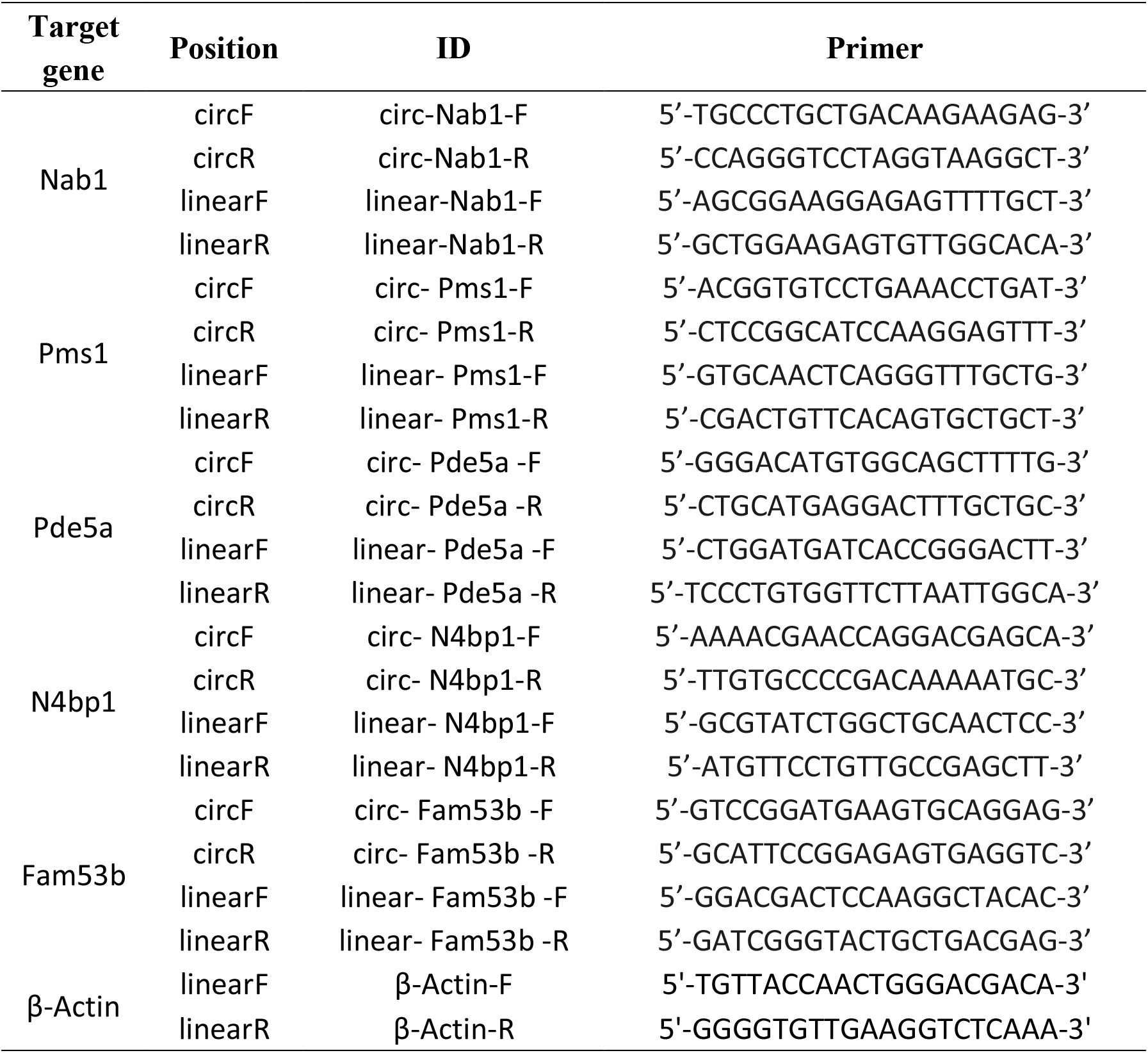
Table of primers to validate differentially expressed circRNAs by RT-qPCR

